# Frequency Conservation Score (FCS): the power of conservation and allele frequency for variant pathogenic prediction

**DOI:** 10.1101/805051

**Authors:** Jose Luis Cabrera Alarcon, Jose Antonio Enriquez, Fátima Sánchez-Cabo

**Author notes:** To whom correspondence should be adressed.

## Abstract

**Background:** Prediction of pathogenic variants is one of the biggest challenges for researchers and clinicians in the time of next-generation sequencing technologies. Stratification of individuals based on truly pathogenic variants might lead to improved, personalized treatments.

**Results:** We present Frequency Conservation Score (FCS) and Frequency Conservation Score for Mitochondrial DNA (FCSMt) two methods for the detection of pathogenic single nucleotide variants in nuclear and mitochondrial DNA, respectively. These scores are based in a random forest model trained over a set of potentially relevant predictors: (i) conservation scores (PhastCons and phyloP); (ii) locus variability at each genomic position built from gnomAD database and (iii) physicochemical distance for amino acids substitutions and the impact/consequence over the canonical transcript. FCS *showed an AUC of 98% for deleteriousness in an independent validation dataset, outperforming other* scores such as m*etaLR, metaSVM, REVEL, DANN, CADD, SIFT, PROVEAN or FATHMM-MKL*. Moreover, FCSMt presented an AUC=0.92 for pathogenic mitochondrial SNVs detection. The tool is available at http://bioinfo.cnic.es/FCS

**Conclusions:** *FCS* and *FCS-Mt* improve pathogenic mutation detection, allowing the prioritization of relevant variants in Whole Exome and Whole Genome Sequencing Analysis.

## 1 BACKGROUND

Most variation between individuals has no direct impact on health. Hence, prioritization of variants according to their potential pathogenicity is a challenge in the detection of genetic based diseases. To help in this task, the American College of Medical Genetics and Genomics (ACMG) and the American Association for Molecular Pathology (AMP) recommended the use of computational prediction tools for the interpretation of the identified variants (1). Therefore, it is clear the need of accurate tools for pathogenic variants detection.

Mendelian diseases are produced mainly by rare or low frequency variants. For this reason, variants detected at low frequency are often classify as potentially pathogenic. However, the definition of “low” frequency relies in an arbitrarily set cutoff. This problem became more apparent when a large number of the variants contained in aggregated databases of population variants, such as ExAc and GNOMAD, were very low frequency single nucleotide variants (SNV) or even singletons (2). Additional to variant frequency, allele frequency could give information for deleteriousness for variant prioritization. In this sense, allele frequency for variants allocated in a specific genetic position, could also give an indirect measure of the relevance of this nucleotide. Bearing in mind the assumption that population variability in a concrete genomic position could be related with the selective pressure associated to this nucleotide, we could assume that the number of variants at a specific position weighted by their frequencies in the population could reflect the relevance of this locus and its pathogenic status. Therefore, allele frequency/locus variability could be a relevant feature to be included in a functional predictor.

The most relevant state of the art tools for the detection of pathogenic variants are: metaSVM, metaLR (3) and REVEL (4). MetaSVM and MetaLR are two ensemble scores based on Support Vector Machine (SVM) and Logistic Regression (LR), respectively. Both methods integrate the information of 11 non-ensemble predictors (PolyPhen-2, SIFT, MutationTaster, Mutation Assessor, FATHMM, LRT, PANTHER, PhD-SNP, SNAP, SNPs&GO and MutPred), three conservation scores (GERP++, SiPhy and PhyloP) and four ensemble scores (CADD, PON-P, KGGSeq and CONDEL) (3). REVEL is also an ensemble score, a random forest algorithm that relies in MutPred, FATHMM v2.3, VEST 3.0, Polyphen-2, SIFT, PROVEAN, MutationAssessor, MutationTaster, LRT, GERP++, SiPhy, phyloP, and phastCons (4). All of them are meta-learners obtained by machine learning, that strongly rely over other functional predictors, outperforming them and proving that machine learning is an interesting strategy to undertake the challenge of pathogenic variants detection because of the large number of variants and samples currently available.

On the other hand, the degree of DNA conservation is also a relevant indicator of nucleotide importance, which could correlate with neutral-pathogenic status. Many of the available tools for deleterious variants detection depend somehow in conservation information, constituting a relevant resource for functional predictors.

Most of these predictors are built to annotate variants encoded in the nuclear DNA. We know, however, that human genetic information is encoded by two widely different genomes, nuclear and mitochondrial genome. Both genomes have their own evolutionary engines: while nuclear genome presents sexual reproduction as source of variability with sister chromatid exchange, mitochondrial DNA is mainly maternally inherited and has a higher mutation rate as its main source of variability. Therefore, mitochondrial DNA has its own conservation path and population frequencies and may not present the same behavior as nuclear DNA for these features. This could be a major point to take into account for the classification of mitochondrial variants.

In this paper we present Frequency-Conservation-Score (FCS) and Frequency-Conservation-Score for Mitochondrial DNA (FCSM), two machine learning methods for the prediction of variant deleteriousness in nuclear and mitochondrial DNA respectively.

FCS and FCSM are freely available at bioinfo.cnic.es/FCS as R shiny app.

## 2 METHODS

FCS and FCSM were built following the work flow depicted in figures 1A and 1B, respectively.

**Fig. 1.**
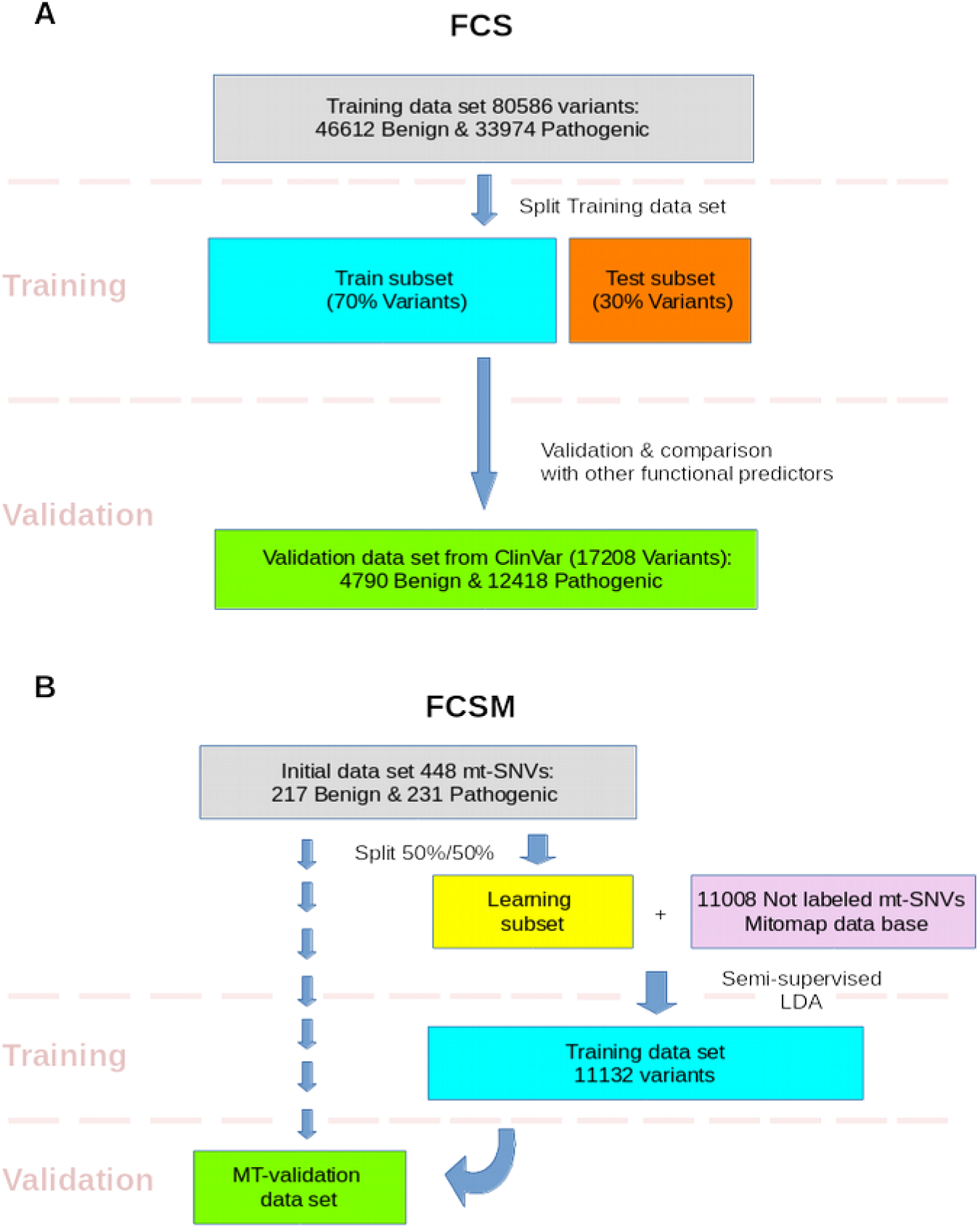
Followed workflow for the development and validation of FCS (A) and FCSM (B). mt-SNVs: mitochondrial single nucleotide variants; LDA: Linear Discriminant Analysis.

### Models training and validation for FCS

We trained four different models, a random forest, a logistic regression, a least absolute shrinkage and selection operator (LASSO) and a neural network. Models specific parameters were tuned up by 5-fold cross-validation, splitting the data into 80% training and 20% evaluation set of **train subset** (supplementary file).

Tuned up models were evaluated in **test subset** and most accurate model measured as area under curve (AUC) in the receiving operator characteristic (ROC) curve was selected as FCS. Then FCS was validated in **ClinVar validation data set**.

Multicollinearity was assessed for explanatory variables in the model (supplementary file). Beside that, variable importance was studied calculating net reclassification improvement NRI and AUC differences (differences in AUC between a model with the variable and a model without the variable within a bootstrap strategy) by calculating a D-statistic, using **ClinVar validation data set**, (supplementary file).

Finally, a cutoff value for FCS score was proposed as a trade between sensibility and specificity, for pathogenic variant detection.

**In brief, for FCS development the score followed a double validation, first in test subset where FCS was selected and second in ClinVar validation data set.**

Models were trained using caret v-6.0 (5), glmnet v-2.0 (6), ranger v-0.11.2 (7) and nnet v-7.3 (8) R-packages. Received Operative Curves and their respective Areas Under the Curve were obtained by pROC v-1.15.0 (9) and ROCR v-1.0 (10) R-packages. NRI was calculation PredictABLE v-1.2.2 R-package (11) and D statistic was calculated meanwhile pROC.

### Models training and validation for FCSM

The same four models trained for FCS were also trained for FCSM (random forest, logistic regression, LASSO and neural network), tuning up their parameters by 5-fold CV. Then, obtained models from training step were evaluated in **validation data set** in order to select the best model as FCSM, figure 1C and 1D. Multicollinearity was analyzed for variables included in FCSM as well as their relative importance within the model measured as NRI and differences between AUC due to the variable.

A cutoff value for FCSM score was set, as the best trade-off between sensibility and specificity, for pathogenic variant detection in mitochondrial DNA.

**Unlike for FCS, training data set was not split in training and test subset and FCSM was validated only in validation data set.**

### Training data sets

#### Training data set for FCS

The training data set was built gathering unique variants from twelve bench-marking data sets also used for the development of predictors published by other authors (IDSV and MutationTaster2), included in VariBench benchmark database suite (12–14). After filtering variants that were included in validation data set, obtained 80586 variants (46612 benign Vs 33974 pathogenic). Training data set was split into two different subsets, the **training subset**, containing 70% of variants used for training the models and another data subset and the **test subset**, represented by the remaining 30% of variants, used for testing the models in order to select the best model, figure 1A.

#### Training data set for FCSM

To build training data set for FCSM, we gathered 224 variants from high confident Clinvar variants, Mitomap (15) curated variants and Varibench. These labeled variants represent the learning subset, that was used to lead a semi-supervised machine learning strategy, with a Linear Discriminant Analysis (LDA), to assign labels to mitochondrial variants registered for sequences deposited in Genebank, taken from Mitomap. Labeled mitochondrial variants from Mitomap represent the **training data set** for FCSM, figure 1B.

#### FCS and FCSM variables

Considered features to train both FCS and FCSM were the locus variability, phastCons(16) and phyloP(17) conservation scores, Grantham distances and variant’s predicted impact over the canonical transcript.

Locus variability was computed as:

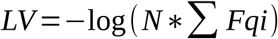

where LV is locus variability, N is the number of alleles described in gnomAD gnomAD 2.1.1 (18) for FCS or in Mitomap data base for FCSM, including the considered variant and Fq_i_ are gnomAD/Mitomap frequencies for alleles affecting to this position. If the variant is not described in the data bases, its frequency is considered to be 0.000001.

The impact over the canonical transcript was obtained using Variant Effect Predictor VEP (19), web version for GRCh37, that classifies it as “HIGH”, “MODERATE”, “MODIFIER” and “LOW”. Variant impact categories were transformed into dummy variables, in order to obtain a coefficient for each category in the regression model, so each category acts as a switcher.

Variants were also annotated with Grantham score for the amino acids substitutions, setting this value to 0 for no missense SNVs (20).

PhastCons and phyloP scores were represented by pre-computed values estimated over a multiple sequence alignment of 100 vertebrate species. AnnotationHub v-2.14.5 (21) and GenomicScores v-1.6.0 (22) R-packages, were used for variant annotation with these conservation scores.

Variants’ data imputation was carried out as mean and mode values, using randomForest R package (23). Before variable imputation the percentages of missing values was a 0% for locus variability, 0% for Grantham score, 0% for each impact dummy variable, 0.65% (585 variants) for Phastcons and 0.65% (585 variants) for phyloP.

### Validation data sets

#### Validation data set for FCS

Validation data set was obtained from variants in ClinVar data base (24,25), selecting variants clinically classified as “benign” or “pathogenic”. These variants were annotated using dbNSFP 4.0a (26), with 3 general ensemble functional predictor scores, MetaLR, MetaSVM, REVEL, CADD (27), DANN (28), SIFT (29), PROVEAN (30) and FATHMM-MKL (31), obtaining 17208 variants (4790 benign Vs 12418 pathogenic).

#### Validation data set for FCSM

Mitochondrial validation data set for FCSM was represented by 224 variants from high confident ClinVar variants, Mitomap curated variants and mitochondrial curated variants from Varibench data sets, other than the variants from learning subset, figure 1B.

#### Comparative study

The accuracy of FCS was compared against other functional predictors, measured as AUC in ROC curves and the performance in Precision-Recall PR curves. For this purpose, it was selected functional predictors commonly used in clinical practice (REVEL, metaSVM, metaLR, CADD, DANN, SIFT, PROVEAN and FATHMM-MKL). Accuracy differences between predictors were evaluated, calculating D statistic score (supplementary file).

For FCSM, the limited amount of pre-computed values for other functional predictors over considered mitochondrial validation data set did not allow FCSM comparison with other predictors. Nevertheless, theperformance of FCSM was compared with FCS in mitochondrial SNVs.

## 3 RESULTS

### FCS RESULTS

#### Selected model

Random forest resulted as the most accurate model in training step showing an accuracy of AUC=0.92, so it was selected as FCS (more details in supplementary file). Regarding to correlation analysis performed, most of model’s regressors showed low correlation level, with the exception of both conservation scores with strong (figure 1 and table 1, supplementary file).

Analyzing variable importance in FCS, measured as NRI values, we obtained that most relevant variable was locus variability NRI=1.4173 [1.3976 - 1.437], p-value<0.001; followed by phyloP score NRI=0.3869 [0.3666-0.4072], p-value<0.001; Phastcons score NRI=-0.0782 [-0.0925--0.0638], p-value<0.001; Grantham’ Score NRI=0.2399 [0.221-0.2588], p-value<0.001; HIGH impact dummy variable with NRI=0.1476 [0.1362-0.159], p-value<0.001; MODERATE impact dummy variable with NRI=0.0694 [0.0597-0.0792], p-value<0.001; MODIFIER impact dummy variable with NRI=0.1259 [0.1149-0.1368], p-value<0.001 and LOW impact dummy variable with NRI=0.1162 [0.1056-0.1268], p-value<0.001 (figure 2 and supplementary file). Although NRI result was negative for PhastCons score, the variable was considered for the model given its D-statistic result D=3.3518 (p-value<0.001).

**Figure 2.**
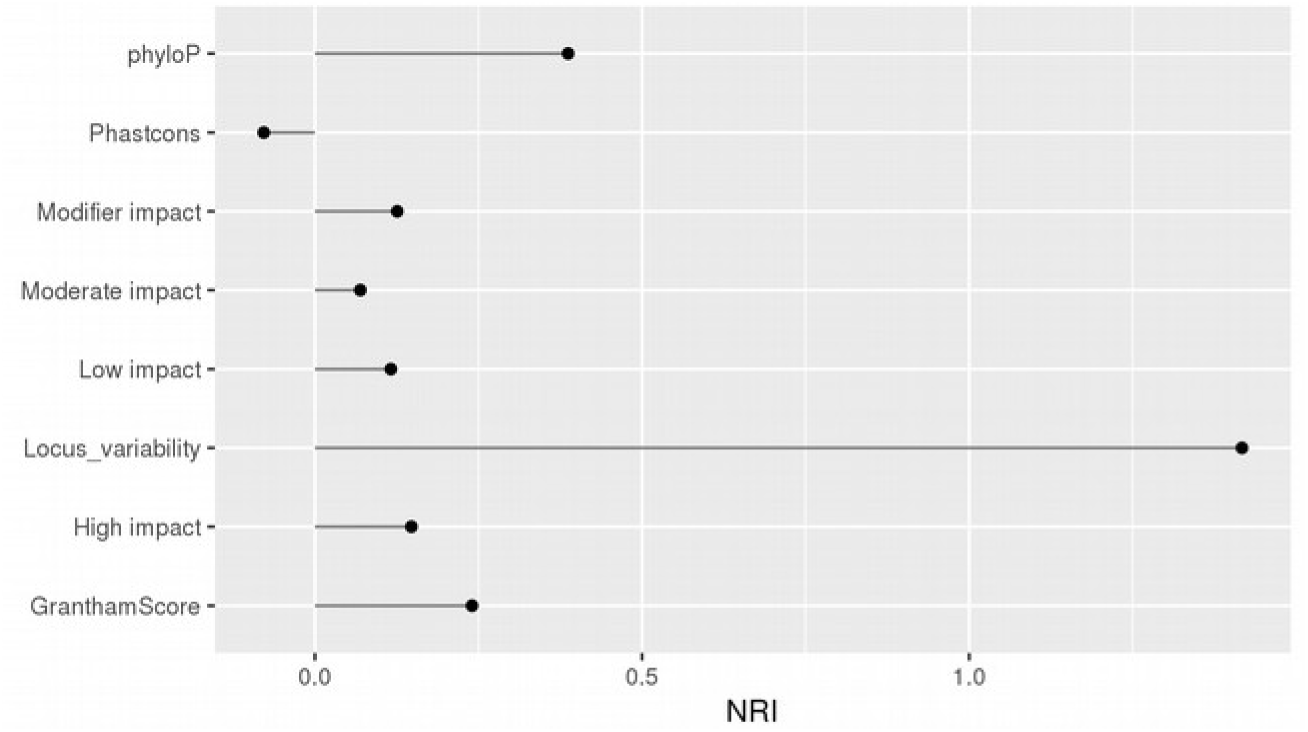
Feature relative importance in FSC, measured as NRI value.

#### AUC comparison

According to our outcomes, FCS showed the highest accuracy AUC=0.98 for pathogenic variant detection and also the highest Precision Recall trade, followed by REVEL (AUC=0.96), metaLR and metaSVM both with AUC=0.93, SIFT and CADD both with AUC=0.90, PROVEAN (AUC=0.89), FATHMM-MKL (AUC=0.84) and DANN (AUC=0.82), figures 3A and 3B. Accuracy differences between scores, computed as D-statistic revealed that FCS was statistically significant better than REVEL (D=13.03; p<0.001), metaLR (D=21.893; p<0.001), metaSVM (D=24.553; p<0.001), CADD (28.736; p<0.001), SIFT (D=29.019; p<0.001), PROVEAN (D=30.864; p<0.001), FATHMM-MKL (D=39.151; p<0.001) and DANN (D=41.664; p<0.001).

**Figure 3.**
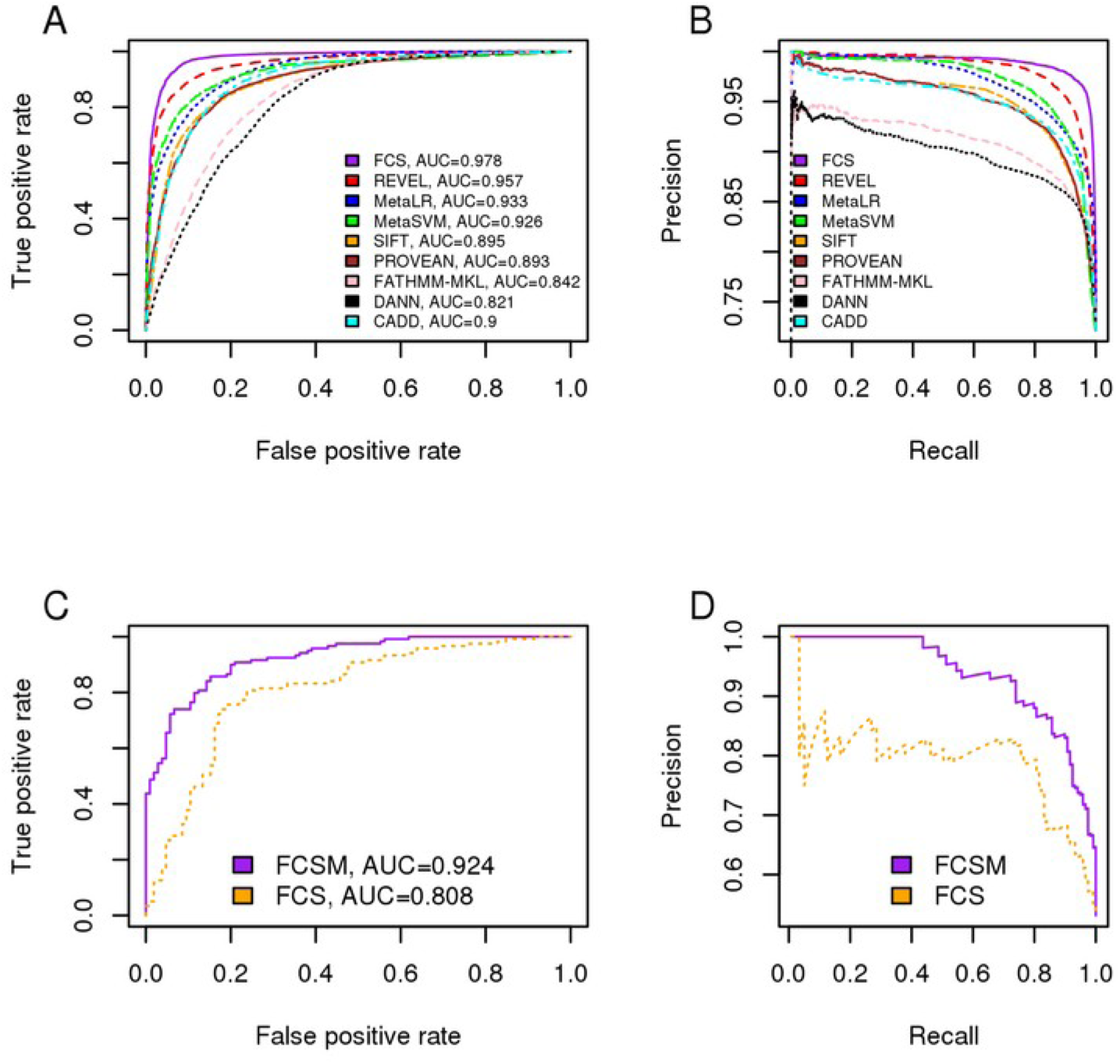
ROC curves (A) and Precision-Recall PR curve (B) for FCS and comparing functional predictors in ClinVar validation data set. ROC curve (C) and PR curve (D) for FCSM in Mitochondrial validation data set.

#### Cutoff value for FCS

According to our validation data, we suggest a cutoff for FCS of 0.4067561, giving 94% of sensibility and 93% of specificity. This threshold was selected as the lowest value of FCS with the best trade-off sensitivity/specificity.

### FCSM RESULTS

#### Selected model

Random forest presented the highest accuracy AUC=0.92, followed by LR model AUC=0.87, LASSO AUC=0.81 and the neural network AUC=0.5 (figure 2 supplementary file). Therefore, Random forest model was selected as FCSM, figures 3C and 3D. According to features relative importance analysis, Locus variability presented a NRI=1.1154 [0.9136-1.3172], but none of the other variables presented a significant evidence (NRI or D-statistic values) for feature relative importance (table 3 supplementary file). Bivariate association study and correlation analysis between features included in FCSM reveled that most of the variables showed low association degree, figure 2 supplementary file.

#### FCS Vs FCSM comparison for mt-SNVs

FCSM (AUC=0.92) outperformed FCS (AUC=0.81) for neutral-pathogenic classification of SNVs in mitochondrial DNA, both in terms of accuracy and precision-recall trade-off, figure 3C and 3D.

#### Cutoff value for FCSM

Considering our outcomes the best threshold of FCSM for pathogenic variant detection in mt-DNA was 0.488 rendering 0.86% of sensibility and 0.85% of specificity.

## 4 DISCUSSION

We have developed FCS and FCSM, two methods to discriminate neutral from deleterious SNVs, in nuclear DNA and mitochondrial DNA respectively. Regarding to ROC curves comparison results, FCS reached the highest accuracy compared to the other considered scores, that are widely used as predictors for variant pathogenicity. Being a not stacked machine learning score, FCS uses information resources that could represent an added value in variant ranking. REVEL, metaLR and MetaSVM, are three of the most accurate predictors published in the literature in pathogenic variant detection. All of them are machine learning based approaches that share most of their constituent features, independently of trained underlying algorithm. In this sense, FCS only shares with them the use of conservation scores, but also includes additional information, as the locus variability derived from gnomAD, the physicochemical impact in amino acids substitutions gathered up in Grantham score and the variant impact over considered canonical transcript, allowing to improve other scores results in variants pathogenic-neutral classification. However, this increased accuracy was joined to the best performance in PR curves, so FCS presented the best results with the least costs in terms of false positives and false negatives.

Though nuclear and mitochondrial DNA share a co-evolution track, for SNVs classification in mt-DNA, it is necessary to take into account, that their different evolving strategies lead to differences in locus variability and conservation status. Therefore, FCSM trained over the same regressors as FCS but in mtDNA variants, presented higher neutral-pathogenic classification ability than FCS for SNVs detection in mt-DNA, figures 3C and 3D. The accuracy presented by software predictors over human non-synonymous variants in mtDNA, ranges from 0.48 to 0.84 (32). In this sense, FCSM resulted as a fairly accurate predictor trained for mitochondrial particularities with an AUC=0.92. Additionally, unlike other classifiers, our predictor is trained not only for missense variants affecting proteins, but also for variants affecting tRNA and control region in mitochondrial chromosome, so FCSM could add extra information for variant prioritization in mtDNA.

In this study, unlike the strategy adopted by other authors focusing in allele frequency to select working variants, we decided to include this information to train the random forest algorithm. Nevertheless, allele frequency information was considered only to extract the degree of locus variability, giving an indirect measurement of the relevance and the freedom for diversity at each considered genomic position. In accordance with NRI, locus variability presented the highest relevance in the final outcome, both for FCS and FCSM.

Phastcons and phyloP are two widely used conservation tools, that relies in different strategies (16,17). PhastCons is a hidden Markov model-based method that estimates conservation rate, for a specific site, taking in to account the rates of neighboring sites. By contrast, PhyloP scores measure evolutionary conservation at individual alignment sites, giving information not only about the magnitude but also about the direction of the evolution rate compared with a neutral drift model. The two methods have different strengths and weaknesses, PhastCons is effective for conserved elements/regions detection and phyloP, on the other hand, is more appropriate for evaluating signatures of selection at particular nucleotides or classes of nucleotides. Relaying in different approaches, both scores provide independent and complementary information for FCS, but according to NRI values, Phastcons is more relevant in FCS. On the other hand, there is no evidence for importance interpretation of both scores in FCSM, probably due to validation data set size curse.

Additionally, we also considered the direct effect of SNVs in canonical transcripts as measurable feature to train our models through impact dummies variables. All of them resulted approximately equivalent in FCS, with much lower weight in variant classification than Locus variability, while there were no evidences about their relative importance in FCSM.

Regarding to missense SNVs, Grantham score gives the physicochemical impact underlying in amino acids substitutions, establishing the distance between these amino acids depending on the composition, polarity and molecular volume. This score, though does not take in to account 3D structure of the protein, can place a complementary background to the one given by the conservation scores, focused in evolutionary distances over nucleotide sequence.

Since the development of next generation sequencing technology and its clinical appliance for mendelian diseases diagnostic or cancer management, discriminating deleterious variants from the bast mass of neutral variants, has became a key stone in clinical diagnostic. In this sense, it is capital the use of informative tools that aid in the task of variant prioritization, oriented to reduce the group of variants of uncertain significance. For this purpose, it is important the use of a wide range of information to undertake this task accurately. In this project, we demonstrated that our score FCS, gives a new approach for SNVs pathogenic classification, that improves the performance of other scores commonly used as functional predictors in clinical practice, so could be considered as a tool for variant ranking, except for mitochondrial SNVs, where FCSM has proved to be a better tool.

In future studies, we shall investigate the possibility of improve pathogenic status detection, considering the inclusion of insertion and deletion variants for training a new version of our functional predictor scores.

## 5 CONCLUSIONS

FCS is a tool with a higher accuracy, compared with other relevant scores for pathogenic mutation detection. This improvement may be due to the addition of allele frequency derived information added to the partial detection power given by conservation information, predicted impact over the transcripts or amino acids substitution relative importance. Therefore it could be used to prioritize variants as disease candidates.

FCSM could be used in variant prioritization for SNVs in mt-DNA, given that is a specific score trained considering mt-DNA particularities.

