## Supplementary information for "Frequency Conservation Score (FCS): the power of conservation and allele frequency for variant pathogenic prediction"

**Multicollinearity analysis for FCS and FCSM features**

Multicollinearity may represent a problem in multivariable regression leading to coefficients overestimation of related variables. Therefore, association between variables considered for training the models were studied by bivariate analysis (linear or logistic regression according to compared variables) in order to explore aggressors' relationship, both for FCS and FCSM training data. Additionally, bivariate analysis was complemented computing Spearman correlation scores.

**In FCS training subset**

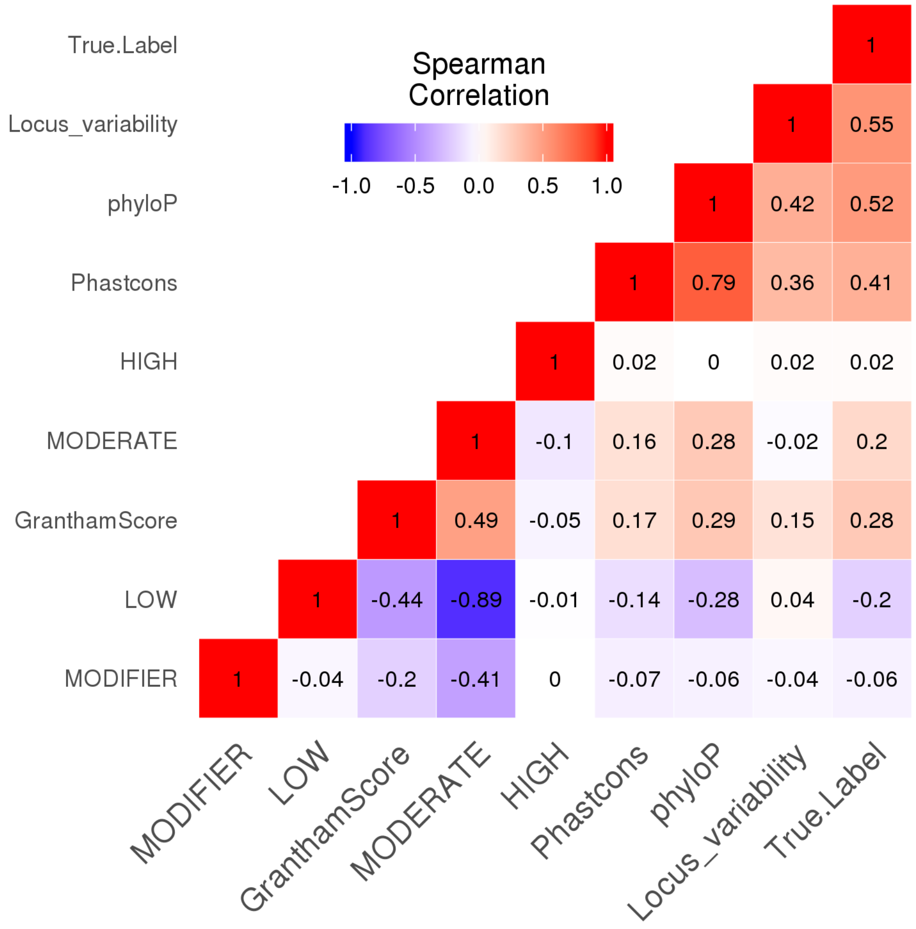


Figure 1. Heatmap for Spearman correlation rho values for FCS model's features.

**Table 1. Bivariate analysis between considered features for FCS in training subset.**

| X / Y | **Locus var** | **PhastCons** | **phyloP** | **Grantham** | **High** | **Moderate** | **Modifier** | **Low** |
|---|---|---|---|---|---|---|---|---|
| **Locus var** | - | β=1.833<br>SE=0.024<br>p<0.001*** | β=0.265<br>SE=0.003<br>p<0.001*** | β=0.006<br>SE=0.0002<br>p<0.001*** | β=1.190<br>SE=0.354<br>p<0.001*** | β=-0.308<br>SE=0.039<br>p<0.001*** | β=-0.667<br>SE=0.087<br>p<0.001*** | β=0.519<br>SE=0.043<br>p<0.001*** |
| **PhastCons** | β=0.052<br>SE=0.0007<br>p<0.001*** | - | β=0.088<br>SE=0.0004<br>p<0.001*** | β=0.0015<br>SE=0.00004<br>p<0.001*** | β=0.223<br>SE=0.060<br>p<0.001*** | β=0.241<br>SE=0.006<br>p<0.001*** | β=-0.261<br>SE=0.015<br>p<0.001*** | β=-0.234<br>SE=0.007<br>p<0.001*** |
| **phyloP** | β=0.513<br>SE=0.005<br>p<0.001*** | β=6.023<br>SE=0.024<br>p<0.001*** | - | β=0.02<br>SE=0.0003<br>p<0.001*** | β=0.139<br>SE=0.492<br>p=0.777 | β=3.53486<br>SE=0.052<br>p<0.001*** | β=-1.559<br>SE=0.121<br>p<0.001*** | β=-3.932<br>SE=0.057<br>p<0.001*** |
| **Grantham** | β=2.345<br>SE=0.081<br>p<0.001*** | β=19.689<br>SE=0.480<br>p<0.001*** | β=3.911<br>SE=0.057<br>p<0.001*** | - | β=-73.8798<br>SE=6.876<br>p<0.001*** | β=81<br>SE=0.672<br>p<0.001*** | β=-2.945<br>SE=34.675<br>p=0.932 | β=-79.523<br>SE=0.761<br>p<0.001*** |
| **High** | β=0.138<br>SE=0.038<br>p<0.001*** | β=1.7503<br>SE=0.520<br>p<0.001*** | β=0.011<br>SE=0.037<br>p=0.777 | β=-2.351<br>SE=35.421<br>p=0.947 | - | β=-20.06<br>SE=577.993<br>p=0.972 | β=-13.65<br>SE=582.971<br>p=0.981 | β=-14.71<br>SE=459.117<br>p=0.974 |
| **Moderate** | β=-0.043<br>SE=0.006<br>p<0.001*** | β=1.120<br>SE=0.031<br>p<0.001*** | β=0.340<br>SE=0.006<br>p<0.001*** | β=7.018<br>SE=38.557<br>p=0.856 | β=-15.900<br>SE=72.195<br>p=0.826 | - | β=-19.092<br>SE=78.897<br>p=0.809 | β=-23.526<br>SE=168.900<br>p=0.889 |
| **Modifier** | β=-0.104<br>SE=0.014<br>p<0.001*** | β=-1.192<br>SE=0.070<br>p<0.001*** | β=-0.122<br>SE=0.010<br>p<0.001*** | β=-0.261<br>SE=0.015<br>p<0.001*** | β=-10.473<br>SE=119.029 p=0.93 | β=-21.075<br>SE=212.632<br>p=0.921 | - | β=-15.548<br>SE=168.9<br>p=0.927 |
| **Low** | β=0.071<br>SE=0.0062<br>p<0.001*** | β=-1.084<br>SE=0.035<br>p<0.001*** | β=-0.403<br>SE=0.007<br>p<0.001*** | β=-3.485<br>SE=31.182<br>p=0.911 | β=-11.0087<br>SE=72.195<br>p=0.879 | β=-23.986<br>SE=212.632<br>p=0.91 | β=-14.026<br>SE=78.897<br>p=0.859 | - |

Features of FCS presented very weak, weak or moderate Spearman correlation values, figure 1. Spearman results suggest a strong negative correlation observed between LOW and MODERATE, but this value may reflect the fact that both are the most represented categories in variant impact and are mutually exclusive. Actually,  Spearman coefficients and bivariate analysis confirmed a strong positive correlation between conservation scores. Additionally, locus variability had a moderate association degree with both conservation scores. On the other hand, locus variability and specially PhastCons were the features that better represented variant impact over canonical transcript.

**In FCSM training data set**

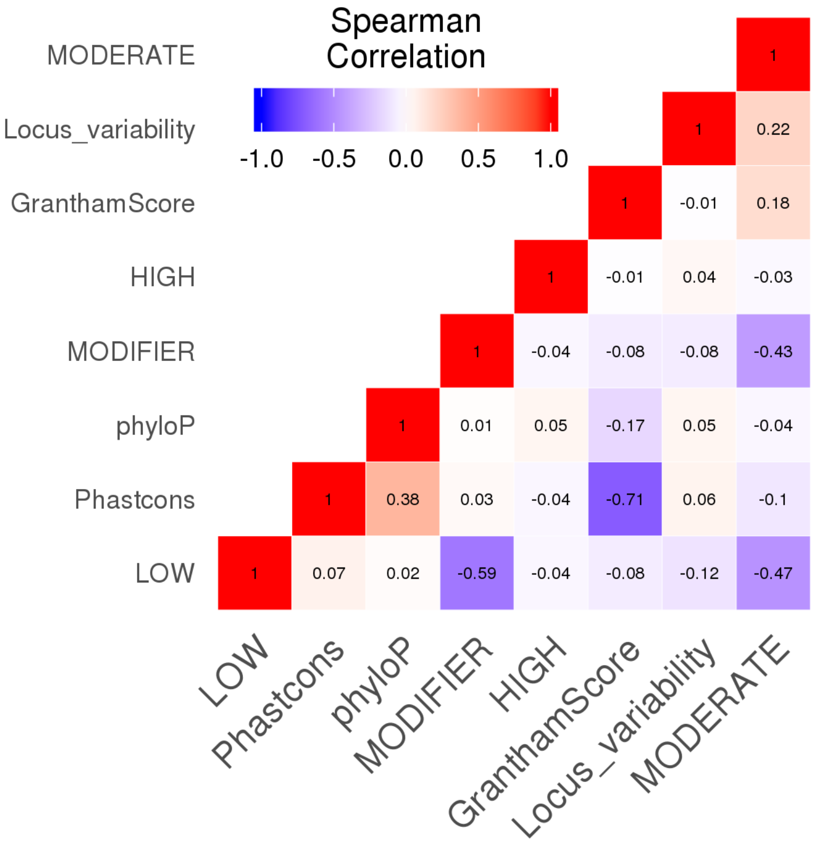


Figure 2.  Heatmap for Spearman correlation rho values for FCSM model's features.

**Table 2. Bivariate analysis between considered features for FCSM in training data set.**

| X<br>Y | Locus var | PhastCons | phyloP | Grantham | High | Moderate | Modifier | Low |
|---|---|---|---|---|---|---|---|---|
| **Locus var** | - | β=0.798<br>SE=0.092<br>p<0.001*** | β=0.055<br>SE=0.012<br>p<0.001*** | β=0.002555<br>SE=0.0012<br>p=0.0312* | β=0.833<br>SE=0.178<br>p<0.001*** | β=0.461<br>SE=0.020<br>p<0.001*** | β=-0.280<br>SE=0.019<br>p<0.001*** | β=-0.11<br>SE=0.018<br>p<0.001*** |
| **PhastCons** | β=0.008<br>SE=0.0009<br>p<0.001*** | - | β=0.064<br>SE=0.001<br>p<0.001*** | β=-0.007<br>SE=0.0001<br>p<0.001*** | β=0.565<br>SE=0.139<br>p<0.001*** | β=-0.021<br>SE=0.002<br>p<0.001*** | β=0.005<br>SE=0.002<br>p=0.0084** | β=0.013<br>SE=0.002<br>p<0.001*** |
| **phyloP** | β=0.034<br>SE=0.007<br>p<0.001*** | β=3.916<br>SE=0.062<br>p<0.001*** | - | β=-0.007<br>SE=0.0009<br>p<0.001*** | β=-0.057<br>SE=0.018<br>p=0.0013** | β=-0.084<br>SE=0.016<br>p<0.001*** | β=0.034<br>SE=0.015<br>p=0.0223* | β=0.029<br>SE=0.014<br>p=0.0447* |
| **Grantham** | β=0.153<br>SE=0.071<br>p=0.0312* | β=-41.040<br>SE=0.604<br>p<0.001*** | β=-0.720<br>SE=0.091<br>p<0.001*** | - | β=-0.675<br>SE=1.379<br>p=0.624 | β=2.629<br>SE=0.157<br>p<0.001*** | β=-1.040<br>SE=0.145<br>p<0.001*** | β=-1.103<br>SE=0.142<br>p<0.001*** |
| **High** | β=1.481<br>SE=0.293<br>p<0.001*** | β=-2.030<br>SE=0.742<br>p=0.0062** | β=0.423<br>SE=0.097<br>p<0.001*** | β=-0.866<br>SE=63.179<br>p=0.989 | - | β=-16.886<br>SE=874.561<br>p=0.985 | β=-17.025<br>SE=745.906<br>p=0.982 | β=-17.084<br>SE=709.534<br>p=0.981 |
| **Moderate** | β= 0.589<br>SE=0.027<br>p<0.001*** | β=-1.834<br>SE=0.202<br>p<0.001*** | β=-0.141<br>SE=0.029<br>p<0.001*** | β=1.231<br>SE=11.587<br>p=0.915 | β=-12.503<br>SE=97.752<br>p=0.898 | - | β=-19.141<br>SE=166.434<br>p=0.908 | β=-19.241<br>SE=158.319<br>p=0.903 |
| **Modifier** | β=-0.291<br>SE=0.020<br>p<0.001*** | β=0.578<br>SE=0.222<br>p=0.0093** | β=0.061<br>SE=0.027<br>p=0.0236* | β=-1.087<br>SE=13.161<br>p=0.934 | β=-12.960<br>SE=97.752<br>p=0.895 | β=-18.460<br>SE=118.359<br>p=0.876 | - | β=-19.875<br>SE=158.319<br>p=0.9 |
| **Low** | β=-0.115<br>SE=0.019<br>p<0.001*** | β=2.114<br>SE=0.358<br>p<0.001*** | β=0.052<br>SE=0.026<br>p=0.0466* | β=-1.101<br>SE=13.0292<br>p=0.933 | β=-13.119<br>SE=97.752<br>p=0.893 | β=-18.659<br>SE=118.359<br>p=0.875 | β=-19.975<br>SE=166.434<br>p=0.904 | - |

Notably, bivariate analysis matched with Spearman correlation results (table 2 and figure 2, respectively). Unlike in FCS training subset (nuclear DNA), conservation scores presented very weak association between them and with locus variability, reflecting an actual difference between both genomes. Moreover, PhyloP showed a weak association with variants' impact, while PhastConst was strong and inversely related to impact over canonical transcript, in an opposite situation to FCS training subset. Focusing on bivariate analysis, locus variability was the feature that better explained variants impact.

**Models training and selection**

We trained four different models: a random forest, a logistic regression, a least absolute shrinkage and selection operator (LASSO) and a neural network, using 5-fold cross validation, splitting the data into 80% training and 20% evaluation set.

Random forest algorithm (both in FCS and FCSM) was trained considering four possible numbers of variables tried at each split, 2, 3, 4 and 5. Tuned up parameters for neural networks were the number of hidden units (from 1 to 10) and the weight of decay that tells how dominant the regularization term will be in the gradient computation (from 0 to 4, by intervals of 0.125).

Models trained were selected according to root mean squared error (RSMSE) in train subset **(in whole training data set for FCSM)**. Selected models were evaluated in test subset **(in validation data set for FCSM)** and most accurate model, measured as the one with the highest area under the receiving operator characteristic (ROC) curve was selected as FCS or FCSM. Finally, FCS was submitted to a second validation using ClinVar validation data set.

Models were trained using caret v-6.0 (McCollum, 2009), glmnet v-2.0 (Friedman, Hastie, & Tibshirani, 2010), ranger v-0.11.2 (Wright & Ziegler, 2017) and nnet v-7.3 (Venables, W. N. & Ripley, 2002) R-packages. For Received Operative Curves performance and comparison of Areas Under the Curve pROC v-1.15.0 (Robin et al., 2011) and ROCR v-1.0 (Sing, Sander, Beerenwinkel, & Lengauer, 2005) R-packages were used.

*Results for FCS:*

The best random forest model, contained 500 trees and 5 variables tried at each split, for selected lasso model the minimum λ=0.00117, tuned up neural network presented an architecture of 10 units in hidden layers and decay=0.125. Random forest (AUC=0.92) outperformed all other trained models in neutral/deleterious variant classification and was selected as FCS, figure 2.

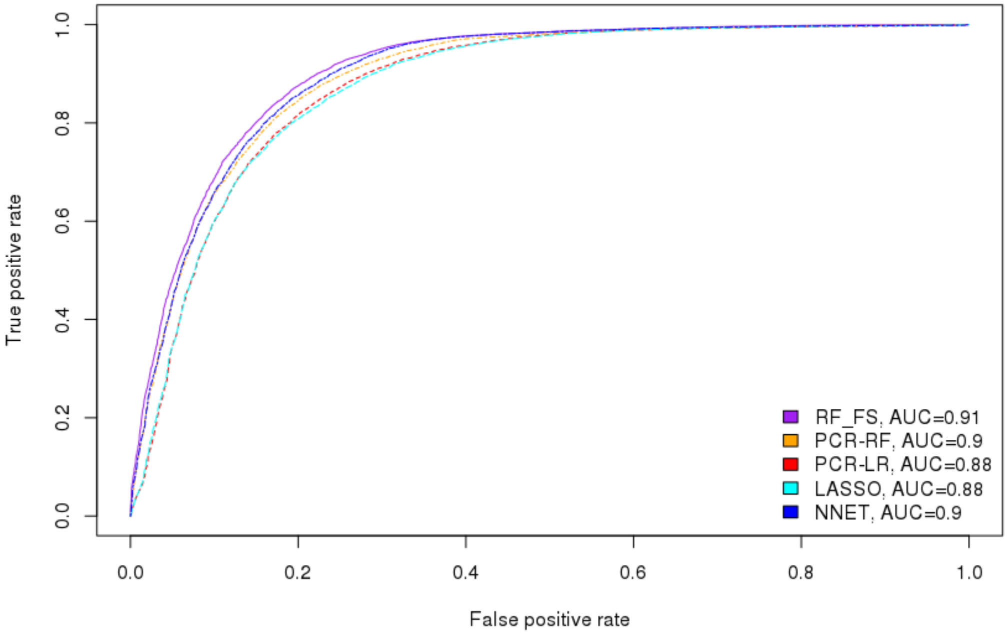


Figure 2. AUC comparison between different trained models in test subset. RF: random forest regression, LR: logistic regression; LASSO: Least absolute shrinkage and selection operator, NNET: Neural network.

*Results for FCSM:*

Selected Random forest model in training step presented 5 variables for splitting at each tree node and 500 trees. Tuned up lasso model had a minimum λ=0.0000613. Selected neural network consisted in a 10 units in hidden layer network and decay=0. As in FCS, random forest (AUC=0.92) outperformed all other trained models in neutral/deleterious variant classification, so was considered as FCSM, figure 3.

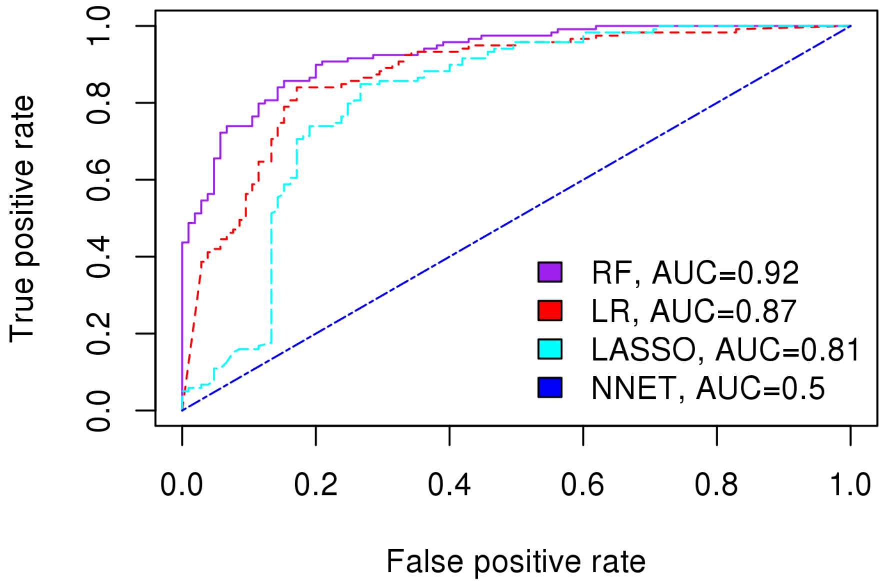


Figure 3. AUC comparison between different trained models for mtDNA. RF: random forest regression, LR: logistic regression; LASSO: Least absolute shrinkage and selection operator, NNET: Neural network.

**Variable importance**

Feature relative importance within selected models was evaluated by computing two different statistics, the Net Reclassification Improvement index and a D statistic computed within 2000 bootstrap re-sampling cycles as:

$$D = \frac{AUC1 - AUC2}{s}$$

where **AUC1** is the area under the curve of predictor number 1, **AUC2** is the area under the curve of predictor number 2 and **s** is the standard deviation for the difference between both values, among n prefixed cycles of bootstrap. We considered 2000 bootstrap cycles. Finally, this D statistic is compared with a normal distribution, to get the probability value. NRI was calculation PredictABLE v-1.2.2 R-package and D statistic was calculated meanwhile pROC R-package. Variable relative importance for FCS and FCSM are gathered in table 3.

**Table 3. Variable importance for FCS and FCSM features measured as NRI and D statistic.**

| | **FCS** | | **FCSM** | |
|---|---|---|---|---|
| ***Feature*** | **NRI [IC 95%] (p-value)** | **D (p-value)** | **NRI [IC 95%] (p-value)** | **D (p-value)** |
| **Locus variability** | 1.4173 [1.3976 – 1.437] (p-value<0.001) | 36.609 (p-value<0.001) | 1.1154 [0.9136 – 1.3172] (p-value<0.001) | 4.0338 (p-value<0.001) |
| **phyloP** | 0.3869 [0.3666 – 0.4072] (p-value<0.001) | 4.9122 (p-value<0.001) | 0.1317 [-0.0267 - 0.29] (p-value=0.1031) | 0.73941 (p-value=0.4597) |
| **PhastCons** | -0.0782 [-0.0925 - -0.0638] (p-value<0.00)1 | 3.3518 (p-value<0.001) | -0.005 [-0.1617 – 0.1516] (p-value=0.94969) | -0.037599 (p-value=0.97) |
| **Grantham Score** | 0.2399 [0.221 – 0.2588] (p-value<0.001) | 3.9407 (p-value<0.001) | 0.0174 [-0.1455 - 0.1802] (p-value=0.83443) | 0.55045 (p-value=0.582) |
| **High impact** | 0.1476 [0.1362 – 0.159] (p-value<0.001) | -2.3809 (p-value=0.017) | 0.0768 [-0.0463 - 0.1998] (p-value=0.22168) | 0.89461 (p-value=0.371) |
| **Moderate impact** | 0.0694 [0.0597 – 0.0792] (p-value<0.001) | -2.92 (p-value=0.004) | 0.0779 [-0.0405 - 0.1963] (p-value=0.1973) | -0.31859 (p-value=0.75) |
| **Modifier impact** | 0.1259 [0.1149 – 0.1368] (p-value<0.001) | -2.5153 (p-value=0.012) | 0.0168 [-0.1009 - 0.1345] (p-value=0.77962) | 0.83945 (p-value=0.4012) |
| **Low impact** | 0.1162 [0.1056 – 0.1268] (p-value<0.001) | -3.0847 (p-value=0.002) | -0.0308 [-0.1623 - 0.1007] (p-value=0.64612 ) | -1.3973 (p-value=0.1623) |

-
